# Mapping Enteric Neural Circuits by Anterograde Transsynaptic Tracing

**DOI:** 10.64898/2026.07.28.741165

**Authors:** Wei Li, Radhashree Sharma, Lei Li, Claire Jyoti Millett, Paul Andrew Muller, Alessandro Furlan, Ulrika Marklund

**Affiliations:** Division of Molecular Neurobiology, Department of Medical Biochemistry and Biophysics, Karolinska Institutet, Stockholm, Sweden; Department of Neuroscience, Karolinska Institutet, Stockholm, Sweden; Kallyope Inc, 430 East 29(th) Street, New York, NY 10016, USA; Perelman School of Medicine at the University of Pennsylvania, Philadelphia, PA 19104, USA

## Abstract

The diverse functions of the enteric nervous system (ENS) arise from communication between molecularly distinct neuronal populations organized into complete circuits. While recent single-cell transcriptomic studies have resolved the molecular identity of enteric neuron classes, methods for defining their synaptic connectivity remain limited. Here, we describe the implementation of mWmC, an anterograde monosynaptic tracer based on a fusion of wheat germ agglutinin (WGA) and mCherry, as a non-toxic, single-component viral tool for mapping neuronal circuits within and beyond the ENS. Following adeno-associated virus (AAV)-mediated expression in enteric neurons, mWmC was efficiently expressed and transmitted selectively to postsynaptic neurons, with no detectable transfer to enteric glia, interstitial cells of Cajal, blood vessels or other mesenchymal cell types. The method also identified postsynaptic neurons in the celiac-superior mesenteric ganglia following tracing of intestinofugal enteric neurons, demonstrating its utility for mapping inter-organ circuits. As proof of principle, we applied mWmC to two genetically defined myenteric interneuron populations and identified preferential postsynaptic targets, revealing selective connectivity with distinct enteric neuron classes. Time-course experiments showed that transsynaptic labeling occurred between 4 and 10 days and reached a plateau thereafter, consistent with monosynaptic transfer. Finally, we developed a dual-reporter version of the system that simultaneously distinguishes input and target neurons within the same tissue. Together, mWmC provides a robust approach for defining circuit architecture in the ENS, linking molecular cell atlases with neuronal connectivity and paving the way for deeper insights into the circuit mechanisms underlying gut physiology.

**Graphical Abstract:** Schematics illustrating the implementation and applications of the anterograde monosynaptic tracer mWmC for mapping enteric neuronal circuits. Parts of schematics are generated with Biorender.

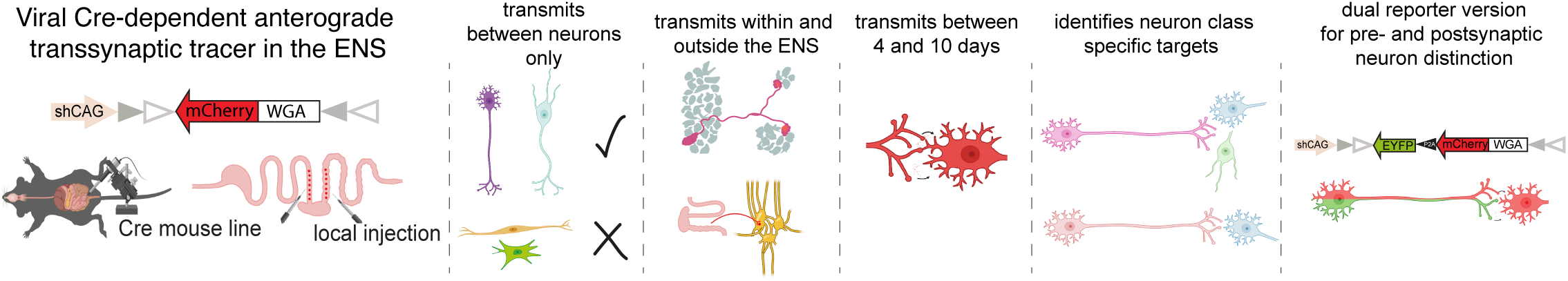

## Main Text

The enteric nervous system (ENS) provides the gastrointestinal (GI) tract autonomous control of essential GI functions through its interconnected neuronal cell types.^1^ Recent single cell RNA sequencing (scRNA-seq) studies have generated unprecedented resolution in the classification of molecular subtypes within the ENS.^2,3^ Yet, methods defining how each neuronal class connects within circuits remain limited. Expression of fluorescent pre-synaptic proteins in specific neuron classes combined with immunostaining of putative target cells has revealed potential communication patterns,^3,4^ but these approaches do not address synaptic transmission. This technical advance introduces an anterograde transsynaptic tracing system to map ENS circuits based on wheat germ agglutinin (WGA) (Graphical Abstract). In the brain, WGA has been demonstrated to travel from soma to axons, released through pre-synaptic terminals and uptaken into post-synaptic neurons.^5^ Recent advances have improved WGA-based transsynaptic tracing through cell-type specific expression of WGA by specific promoters or Cre-mice,^6,7^ and fusion of WGA to mCherry (referred to as mWmC), enabling easier visualization and restricting transmission to first-order synaptic partners.^8^ Here, we describe the implementation of mWmC in the ENS for the study of neuron-class specific circuits. First, we investigated mWmC expression within the ENS and its transmission selectivity. AAV-DIO-mWmC was injected into the distal ileum of pan-neuronal Baf53b-Cre mice and tissues were harvested 4 weeks post-transduction (WPT) (Figure 1A; Supplementary Figure 1). mWmC was detected in HuC/D+ neurons in both the submucosal and myenteric plexuses. In contrast, SOX10+ glia cells were not labelled, thus mWmC does not transmit to enteric glia despite reported nerve varicosities on glial cells^1^ (Figure 1B-E). Because the ENS regulates vascular tone^1^, we examined potential transmission of mWmC to blood vessels but detected no mWmC signal along CD31+ vessels (Figure 1F). A few MHCII+ muscular macrophages stained for mWmC, likely resulting from synaptic pruning or other phagocytic activity (Figure 1G). To specifically target NOS1+ inhibitory motor neurons and addressing potential transmission to interstitial cells of Cajal (ICC), we injected AAV-DIO-mWmC into Nos1-Cre mice and analysed at 4 WPT. We stained for the ICC marker c-KIT and focused on the circular muscle plane guided by mWmC+ projections from the myenteric plexus. No overlap was observed between c-KIT and mWmC, while all mWmC+ cells expressed HuC/D (Figure 1H-J). Overall, we show that mWmC can be readily expressed in enteric neurons, but does not transmit to glia, vessels or mesenchymal cells, consistent with observations in the brain^8^.

**Figure 1.**
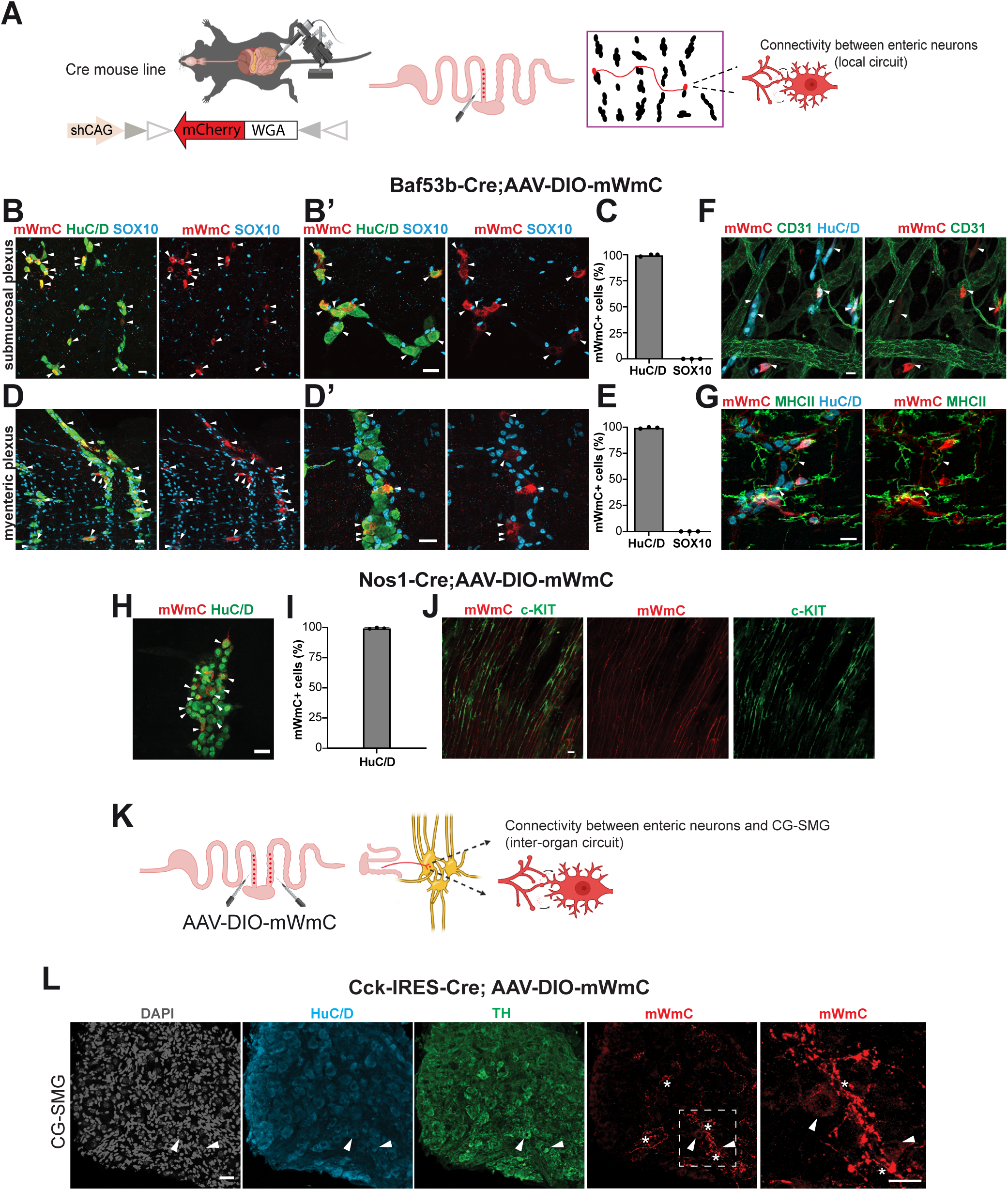
mWmC expressed in enteric neurons transmits to other neurons within and outside the ENS but not to non-neural cells. **(A)** Schematics of local gut injection and AAV construct for Cre-dependent mWmC expression to study postsynaptic transmission in the ENS through anterograde tracing. (**B, D)** Representative confocal images of submucosal plexus (B, B’) or myenteric plexus (D, D’) from ileum of Baf53b-Cre mice injected with AAV-DIO-mWmC and harvested 4 weeks post transduction (WPT), showing mWmC expression in neurons (HuC/D+, arrowheads) but not enteric glia (SOX10+). **(C, E)** Bar graphs showing the percentage of mWmC+ cells that express HuC/D or SOX10 in either the submucosal plexus (C) or the myenteric plexus (E). Each dot represents one mouse. N=3 mice. Data are presented as mean ± standard deviation (S.D). **(F)** Representative picture of the submucosal plexus from ileum of a Baf53b-Cre mouse injected with AAV-DIO-mWmC, showing that mWmC signal (arrowheads) does not overlap with vasculature cells (CD31+). (**G)** Representative picture of the myenteric plexus from ileum of a Baf53b-Cre mouse injected with AAV-DIO-mWmC, showing some mWmC signal in MHCII+ muscular macrophages (arrowheads). (**H)** Representative confocal image of mWmC overlapped with HuC/D+ neurons (arrowheads) in myenteric plexus of a Nos1-Cre mouse analysed at 4 WPT after injection with AAV-DIO-mWmC. (**I)** Bar graph showing the percentage of mWmC+ cells expressing HuC/D. Each dot represents one mouse. N=3 mice **(J)** Representative confocal image displaying the lack of mWmC signal in c-KIT+ ICCs within the circular muscle layer. (**K**) Schematics of local gut injection of AAV-DIO-mWmC to label CCK+ intestinofugal neurons (IFANs) to study postsynaptic transmission in the CG-SMG. (**L**) Representative confocal image of CG-SMG showing extensive axonal networks from mWmC-labelled IFANs (stars) and postsynaptic TH+ neurons (arrowheads) from a Cck-Ires-Cre mouse at 4 WPT after injection with AAV-DIO-mWmC. Right-most picture shows a higher magnification of the area indicated with a white dotted box. Scale bars: 30μm. Parts of (A, K) are created with Biorender.com.

We next asked whether neuronal targets of the ENS outside the gut also can be investigated using mWmC. Intestinofugal CCK+ neurons in the distal ileum and proximal colon project to the Celiac-Superior Mesenteric Ganglia (CG-SMG).^9,10^ To express mWmC in these neurons, we injected AAV-DIO-mWmC into Cck-Ires-Cre mice and collected CG-SMG at 4 WPT. Large numbers of mWmC+ nerve processes were observed, and a few mWmC+ TH+/HuC/D+ neurons were detected, validating the utility of mWmC to study inter-organ circuits (Figure 1K, L).

We next evaluated the potential of mWmC in revealing connectivity patterns between different enteric neuron classes. We have previously defined 12 myenteric neuron classes (myENC1-12)^2^, many of which can be specifically targeted using Cre-mice. Sst-Ires-Cre mice marks myENC5, a potential interneuron class for which the connectivity is little known. Sst-Ires-Cre; LSL-h2B-EGFP mice, labeling SST+ neurons with nuclear EGFP, were locally injected with AAV-DIO-mWmC and analyzed at 4 WPT (Figure 2A). Focusing on mWmC+ neurons, we first distinguished input neurons (EGFP+) and target neurons (EGFP-), identifying 53.63±20.17% as target neurons (Figure 2B). Next, we investigated the identity of mWmC+/EGFP- neurons using three markers which together cover the majority of myenteric neuronal classes^2^: NOS1, CALB2 and Neurofilament M (NF-M). Out of total mWmC+/EGFP- neurons, 20.40±5.73% were NF-M+, 46.8±14.9% were CALB2+, while only 3.76±2.51% were NOS1+, indicating a preferential connection to NF-M+ and CALB2+ neurons, but not to NOS1+ neurons (Figure 2C-F). Targets of myENC5 could thus be myENC1-2, myENC6 (CALB2+), myENC7 and myENC12 (NFM+) (Figure 2G). NOS1 does not only mark inhibitory motor neurons but also an interneuron class.^1^ To investigate targets of NOS1+ interneurons, we analyzed ileum of Nos1-Cre mice injected with AAV-DIO-mWmC at 4 WPT. We found that 55.74±2.78% of mWmC+ neurons lacked NOS1 and thus corresponded to potential targets (Figure 2H, I). We examined CALB2 since it is excluded from NOS1+ neurons and instead labels other classes (myENC1-3; myENC5, myENC6). We found that 21.62±8.25% of mWmC+/NOS1- neurons were CALB2+, indicating robust synaptic connection to CALB2+ neurons (Figure 2I-L). Together, we demonstrate the utility of mWmC in revealing postsynaptic targets of defined ENS classes, exemplified by two interneuron types.

**Figure 2.**
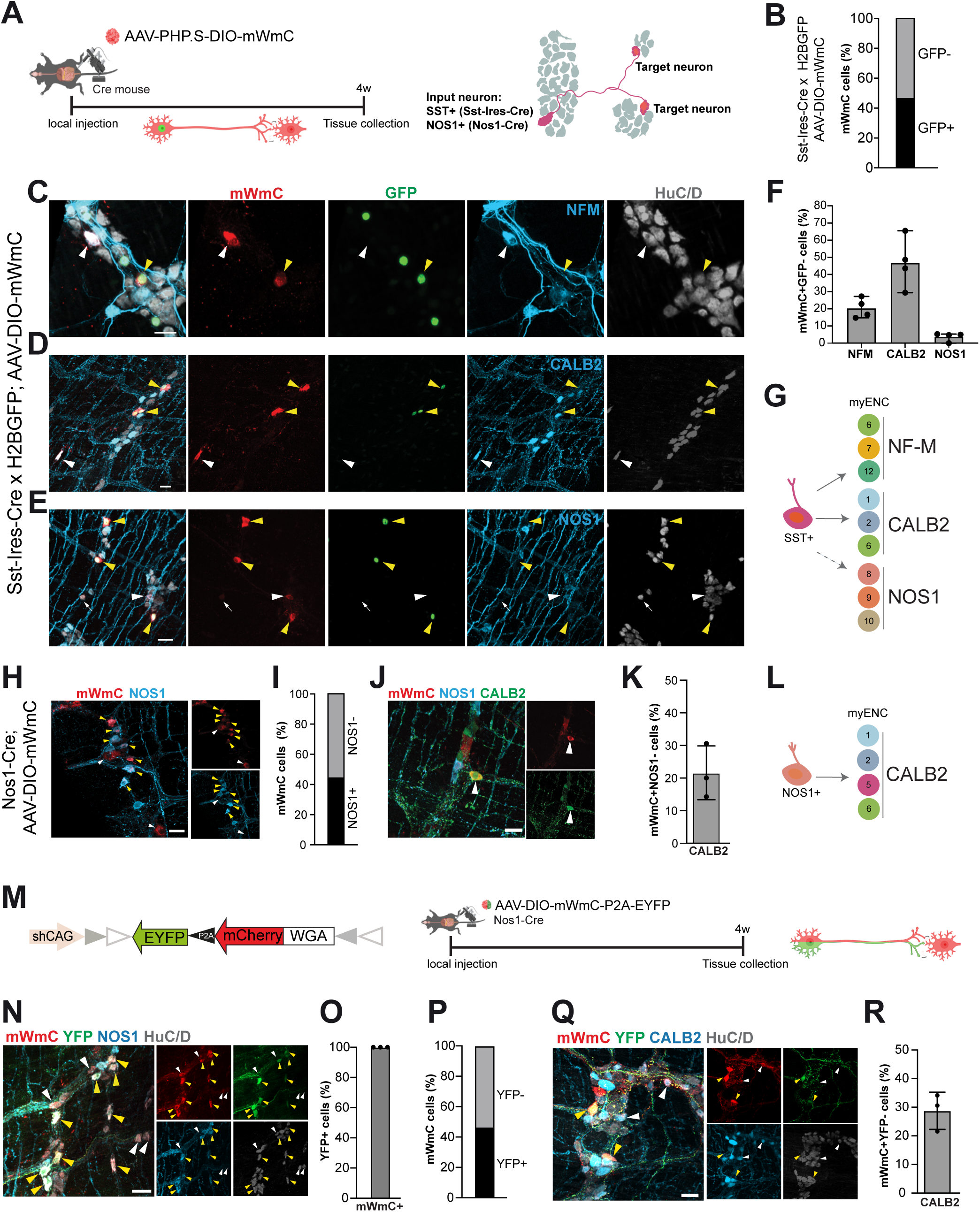
mWmC expression in specific Cre mice reveals neuronal identities of postsynaptic targets of enteric interneuron classes. **(A)** Schematic showing the experimental strategy for investigating postsynaptic neurons of specific interneuron classes using AAV-DIO- mWmC locally injected to the ileum of Cre-mice. (**B)** Stacked bar graph displaying the percentage of mWmC+ neurons that are GFP+ or GFP-. In total 868 mWmC+ cells were counted. N=4 mice. (**C-E)** Representative confocal images of mWmC+ neurons in Sst-IRES-Cre x LSL-h2b-GFP mice. Yellow arrowheads indicate mWmC+/GFP+ neurons (input neurons). White arrowheads in (C, D) indicate mWmC+/GFP- neurons (target neurons) co-stained with subclass markers. White arrow in (E) indicates mWmC+/GFP- neuron that is not positive for NOS1. (**F)** Bar graphs showing the percentage of mWmC+/GFP- neurons that were positive for either subclass marker. Each dot represents one mouse. N=4 mice. Data are presented as mean ± S.D. (**G)** Schematic summary of the preferential connectivity pattern of SST+ neurons. Note that even though myENC5 also expresses CALB2, this neuron class could not be evaluated as a potential target neuron as it cannot be distinguished from input neurons. (**H)** Representative pictures displaying NOS1+ (yellow arrowhead; input neurons) and NOS1- neurons (white arrowhead; target neurons) labeled by mWmC. (**I)** Stacked bar graph showing the ratio between NOS1+ and NOS1- neurons that showed mWmC signal. In total 456 mWmC+ cells were counted. N=3 mice. (**J)** Representative picture of a target mWmC+/CALB2+ neuron (white arrowhead). (**K)** Bar graph displaying the percentage of mWmC+/NOS1- neurons expressing CALB2. Each dot represents one mouse. N=3 mice. Data are presented as mean ± S.D. (**L)** Schematic summary of the connectivity of NOS1+ neurons to CALB2+ neurons (potential myENC1, 2, 5 and 6). (**M)** Schematics showing the dual-reporter construct and the experimental setup for the simultaneous expression of mWmC and EYFP using this construct in the Nos-Cre mouse line. (**N)** Representative confocal image showing mWmC+/YFP+/NOS1+ input neurons (yellow arrowhead), and mWmC+/YFP-target neurons (white arrowhead). **(O)** Bar graph displaying the percentage of YFP+ neurons that express mWmC. N=3 mice **(P)** Stacked bar graph showing the ratio between YFP+/mWmC+ (input) and YFP-/mWmC+ (target) neurons. In total 616 mWmC+ neurons were counted. N=3 mice. **(Q)** Representative confocal image showing mWmC+/YFP+ input neurons (yellow arrowhead) and mWmC+/YFP- target neurons that are also CALB2+ (white arrowhead). **(R)** Bar graph showing the percentage of YFP-/mWmC+ neurons expressing CALB2. Each dot represents one mouse. N=3 mice. Data are presented as mean ± S.D. Scale bars: 30µm. Parts of (A, M’) are created with Biorender.com.

We next addressed the time course over which mWmC transmits from input to target neurons by delivering AAV-DIO-mWmC into the ileum of Sst-Ires-Cre; LSL-h2B-EGFP mice and examining the percentage of GFP-/mWmC+ neurons at 2, 4, 10 and 28 days post transduction (DPT) (Supplementary Figure 2A). We found comparable low levels at 2 and 4 DPT, while the percentage of GFP-/mWmC+ increased to 49.96±7.06% at 10 DPT and 53.63±20.17% at 28 DPT (Supplementary Figure 2B-F). Thus, most anterograde transmission occurred between 4 and 10 DPT, while the similar labeling at 10 and 28 DPT supports the monosynaptic nature of mWmC.

Lastly, we designed and implemented a dual-reporter construct (AAV-DIO-mWmC-P2A-EYFP) to visualize and discriminate input neurons (EYFP+/mWmC+) and target neurons (EYFP-/mWmC+) within the same recombinase system (Figure 2M). We injected this construct into Nos1-Cre mice and found that all EYFP+ cells expressed mWmC, validating the construct fidelity (Figure 2N, O). The ratio of input-to-target neurons, and the ratio of CALB2 target neurons were comparable to the findings using the AAV-DIO-mWmC single-reporter construct (Figure 2N, P-R). EYFP+/mWmC+ boutons adjacent to CALB2+ target cells (Figure 2Q, white arrows) could constitute possible transmission sites.

Altogether, we describe the implementation of the anterograde monosynaptic tracer mWmC, a non-toxic, single-component viral tool for mapping the connectivity of genetically defined neurons within and outside the ENS. Using this method (see Supplementary Material for details), we begin to identify preferential neuronal targets of two myenteric interneuron types. mWmC provides a new approach for linking molecular ENS cell atlases to circuit architecture, paving the way for deeper insights into the circuit mechanisms underlying gut physiology.

## Acknowledgements

We sincerely thank Dr Xin Duan for sharing the DIO-mWmC vector construct and for insightful discussions. We thank the Viral Vector Facility of the Neuroscience Center Zurich for excellent service, plasmid cloning and production of AAV. We thank Prof. Castelo-Branco for sharing LSL-h2b-GFP mice. We thank Linnéa Backman and animal staff from the Department of Comparative Medicine for help with animals and the Biomedicum Imaging Core for excellent Imaging facilities. We acknowledge all members of the Marklund lab as well as Dr. David Linden, Dr. Arthur Beyder and Dr. Kristen Smith-Edwards (Mayo Clinic, Rochester, US) for constructive discussions. We thank Dr. Michael Fatt (KI, Sweden) and Müge Altinkök (KI, Sweden) for constructive feedback of their use of WGA in the brain.

