## Supplementary Material for "Mapping Enteric Neural Circuits by Anterograde Transsynaptic Tracing"

**Supplemental Information**

**Material and Methods:**

**Mice**

The generation of the Baf53b-Cre[1–3] (JAX, #027826), Sst-Ires-Cre[1,4] (JAX, #013044), Nos1-Cre[1,5] (JAX, #017526), Cck-Ires-Cre[2] (JAX, #012706) and LSL-h2b-GFP[6] (JAX #036761) mouse strains have been previously described. Animals were group-housed, with food and water *ad libitum,* under 12-h light-dark cycle conditions, 22°C ambient temperature and 50% humidity. Animal experiments were approved by the local ethics committee in northern Stockholm (Stockholm Norra djurförsöksetiska nämnd, Jordbruksverket; N5237-2023; 1613-2024).

**Viral constructs**

ssAAV-PHP.S/2-CAG-dlox-WGA2_mCherry-dlox-WPRE-hGHp(A) (AAV-DIO-mWmC), and ssAAV-PHP.S/2-shortCAG-dlox-WGA2_mCherry-P2A-EYFP-dlox-WPRE-hGHp(A) (AAV-DIO-mWmC-P2A-EYFP; #v1263) were constructed at the Viral Vector Facility (VVF) of the Neuroscience Center Zurich (ZNZ). AAV-DIO-mWmC was produced using the plasmid *pssAAV-2-CAG-dlox-lgK_3xHA_WGA2_mCherry(rev)-dlox-WPRE(mut6A)-hGHp(A)* (#p909). AAV-DIO-mWmC-P2A-EYFP was produced using plasmid *pssAAV-2-hSyn1-chI-hChR2(H134R)_2A_EYFP_3xNLS-WPR#-bGHp(A)* (#p1182) and *pssAAV-2-CAG-dlox-lgK_3xHA_WGA2_mCherry(rev)-dlox-WPRE-hGHp(A)* (#p1002). Plasmid p909 was a kind gift from Dr. Xin Duan. Viral titers used: AAV-DIO-mWmC: 3.3*10^12 vg/ml; AAV-DIO-mWmC-P2A-EYFP: 5.8* 10^12 vg/ml.

**Local injection to the mouse gut**

Representative pictures of the set-up and key procedures can be viewed in Supplementary Figure 1A-E. Surgical instruments were sterilized using a glass bead sterilizer. Mice were anesthetized in an induction chamber with 3% isoflurane (Attane Vet) in oxygen and maintained under anesthesia with 1.5% isoflurane delivered through a nose cone. Preoperative medication included subcutaneous administration of carprofen (Rimadyl) diluted in saline and Ophthalmic ointment (Viscotears®) was applied to the eyes to prevent corneal drying. The abdomen area was shaved using an Isis rodent shaver (Aesculap), and hair removal cream (Veet) was applied to remove the abdominal hair. The mice were positioned on a sterile surgical drape (Baxter) placed over a heating pad and stabilized by taping four limbs onto the drape. The surgical site was disinfected sequentially with povidone-iodine and alcohol three times. A vial containing AAV was thawed on ice and the virus was loaded into a glass micropipette from 2µl of AAV that had been pipetted on a piece of parafilm. Fast green (Sigma-Aldrich) was added at 0.1% to the virus solution to visualize the injection site. An incision was made at the midline of the abdominal wall to expose the peritoneal cavity, following the confirmation of loss of recoil paw compression. A sterile drape with a central opening was placed over the rest of the mouse body, exposing only the surgical field (Supplementary Figure 1B, D). The distal ileum was located based on its position relative to the cecum and placed on sterile gauge saturated with saline for injection. All injections were made with a custom beveled glass pipette (Clunbury Scientific: using Drummond Cat. No. 3-000-203-G/XL, 7´´ glass, with the parameters: outer diameter: 1.14mm; length: 3.5 inch; inner diameter: 0.53mm; tip outer diameter: 27µm) mounted on a Nanoinjector (NANOLITER2020, WPI). In total 2µl virus was delivered into the ileum wall, with 50nl in each injection site at 15nl/second. 10-15 injection sites were distributed along one side of the intestine. The intestine was thereafter rotated, and another 10-15 injections were made on the opposite side. For trans-organ tracing, 10-15 additional injections were made into the proximal colon on each side (Supplementary Figure 1D). Following each injection, the needle was left in place for 1-2 seconds to minimize reflux. Any unsuccessful injections were immediately cleaned with sterile cotton swap. After completing the injection, the abdominal musculature was closed with absorbable sutures (Ethicon), and the skin was closed with suture clips (Agnthos, #12020-00) and xylocaine ointment (Aspen) was applied on the abdomen to provide local analgesia and reduce irritation. The animal was monitored on a recovery pad until fully awake and mobile. Once fully recovered, the mouse was returned to its home cage. Postoperative care included carprofen administration and daily monitoring of the incision site, defecation, and general health status.

**Tissue preparation for histochemical analysis**

*Intestine:* Ileal segments from injected mice were dissected and placed into PBS on ice. The mesentery was removed and the ileum was opened lengthwise along the mesenteric border. The intestinal contents were rinsed out with ice-cold PBS. The tissue was stretched and pinned with the mucosa side down on a Sylgard (Dow)-coated dissection dish with DMEM. Longitudinal muscle-myenteric plexus-circular muscle layer was peeled off (myenteric peels). Myenteric peels were stretched on a Sylgard-coated dissection dish and fixed in 4% paraformaldehyde (PFA) at 4°C overnight.

*Celiac-Superior Mesenteric Ganglia (CG-SMG)*: The mouse was placed in the supine position, and a midline incision was made through the abdominal skin and peritoneal wall to expose the abdominal viscera. The visceral organs were gently exteriorized using cotton swabs to expose the left kidney. The CG-SMG complex was identified at the junction of the descending aorta and the left renal artery. Using fine forceps and microdissection scissors, CG and SMG were dissected. The ganglia were fixed overnight in 4% PFA, washed twice in PBS, and cryoprotected in 30% sucrose. Samples were then embedded in Neg-50 embedding medium

(Epredia, Cat. No. 6502) and cryosectioned at a thickness of 20 µm using a cryostat (HM 560 Microm International GmbH, Germany).

**Immunohistochemical analysis**

Myenteric and submucosal peels were washed three times with PBS, and cut into 1 $cm^2$ segments and incubated in blocking solution containing purified donkey anti-mouse Fab (Jackson Laboratories, 715-007-003) diluted 1:50 in PBS and 0.3% Triton X-100 (Sigma-Aldrich) overnight at 4°C. The blocking solution was replaced by incubation buffer containing 2% normal donkey serum (NDS; Jackson Laboratories), 1% bovine serum albumin (BSA) and 0.3% Triton X-100 in PBS for 8 hours (over day) at 4°C. Peels were transferred to 48-well plates and incubated in primary anti- and nano-bodies diluted in the incubation buffer for 48 hours at 4°C, washed three times with PBS (10 minutes each) and then placed in the secondary antibodies diluted in the incubation buffer for 2 hours at room temperature in dark with shaking. The tissue was washed three times with PBS (15 minutes each) and mounted on glass slides using DAKO mounting medium (Agilent) containing DAPI. CG-SMG slides were washed in PBS and incubated in blocking solution for 2 hours. Blocking solution was replaced with primary antibodies diluted in incubation buffer over night at 4°C. Slides were washed three times (10 minutes each) and incubated for 1 hour with secondary antibodies diluted in incubation buffer. Slides were washed three times (10 minutes each) and mounted using DAKO mounting medium containing DAPI.

Primary Antibodies:

| **Target:** | **Host:** | **Source:** | **Dilution:** |
|---|---|---|---|
| CD31 | Goat | Novus Bio AF3628 | 1:500 |
| Calretinin | Chicken | EnCor Bio; CPCA-Calret | 1:2000 |
| GFP | Sheep | Biogenesis | 1:1000 |
| GFP | Chicken | Abcam ab13970 | 1:1000 |

| | | | |
|---|---|---|---|
| HuC/D | Human | A gift from David Linden and Vanda Lennon (Mayo Clinic Rochester, US) | 1:20 000 |
| NF-M | Mouse IgG2a | Abcam ab7794 | 1:500 |
| NOS1 | Goat | Abcam ab1376 | 1:1000 |
| RFP/Tomato | Rabbit | Rockland | 1:1000 |
| SOX10 | Rabbit | Abcam | 1:200 |
| TH | Sheep | Novus Bio NB300-110 | 1:300 |

Secondary Antibodies:

| **Antibodies:** | **Source:** | **Dilution:** |
|---|---|---|
| Donkey anti-chicken 488 | Jackson ImmunoResearch 703-545-155 | 1:400 |
| Donkey anti-goat 488 | ThermoFisher A11055 | 1:400 |
| Donkey anti-goat 555 | ThermoFisher A21432 | 1:1000 |
| Donkey anti-goat 647 | ThermoFisher A21447 | 1:250 |
| Goat anti-mouse IgG2a 647 | ThermoFisher A21241 | 1:250 |
| Donkey anti-rabbit 647 | ThermoFisher A31573 | 1:250 |
| Donkey anti-rabbit 555 | ThermoFisher A31572 | 1:1000 |
| Donkey anti-sheep 488 | ThermoFisher A11014 | 1:400 |
| Donkey anti-Human 647 | Jackson ImmunoResearch 709-175-149 | 1:250 |
| Donkey anti-Human 405 | Jackson ImmunoResearch 709-475-149 | 1:200 |
| Donkey anti-Human 488 | Jackson ImmunoResearch 709-545-149 | 1:400 |

**Confocal Imaging**

Imaging of z-stacks with interval of 1μm was performed using a Zeiss LSM980 confocal microscope (Oberkochen, Germany) using 20X and 40X objectives. The images were then processed in Fiji image analysis software (version 2.0.0-rc-69/1.52i) (National Institutes of Health, Bethesda, MD). Color was in some cases changed and each channel was individually adjusted.

**Cell counting and statistical analysis**

Counting of fluorescent cells was performed on confocal images. GraphPad Prism (version 10) was used to generate bar plots. One-way ANOVA with Tukey's multiple-comparison test were used to determine the statistical significance in Supplementary Figure 2B.

**Marker selection and neuron nomenclature**

In this study we have adopted the nomenclature of molecular myenteric neuron classes of Morarach et al (2021),[2] myENC1-12, as defined by single cell RNA-sequencing complemented with immunohistochemical marker validation. Calretinin (CALB2, *Calb2*) marks myENC1-2 (putative excitatory motor neurons), myENC5 (putative interneuron), myENC6 (Intrinsic Primary Afferent Neuron, IPAN). Nitric Oxide Synthase 1 (NOS1, *Nos1*) marks myENC8-9 (inhibitory motor neurons) and myENC10 (putative interneurons). Neurofilament M (NF-M; *Nefm*) marks primarily myENC12 (putative interneurons and mechanosensitive IPANs), but also the majority of myENC7 (putative interneurons and IFAN) and myENC6 (IPAN).

**Comparison to alternative synaptic tracer systems**

Various tracing tools including trans-synaptic chemicals, proteins and neurotropic viruses have been widely used in the central nervous system (CNS).[7,8] Genetically modified rabies virus was developed to infect specific neuron types that express Avian sarcoma leucosis virus receptor, TVA and glycoprotein. Once inside these target neurons, the virus spreads retrogradely across synapses to presynaptic input neurons. Because the input neurons lack the glycoprotein, further transmission is blocked allowing precise circuit mapping.[9,10] However, it requires two separate viral components and injections, adding complexity to the experimental design. In addition, rabies virus is cytotoxic and leads to neuronal loss after extended incubation. Another tool that is recently adopted in the CNS field is Adeno-associated virus serotype 1 (AAV1) that has been shown to act as an anterograde transsynaptic tracer. However, AAV1 as a tracer cannot mediate tracing from genetically defined neurons and is only suitable for studying connectivity between separated anatomical regions.[11,12] In addition, AAV1 exhibits low transduction efficiency in the ENS.[13] In comparison to these methods, the mWmC system may currently allow the most flexible transsynaptic anterograde approach.
