## Supplementary Figure 1 for "Mapping Enteric Neural Circuits by Anterograde Transsynaptic Tracing"

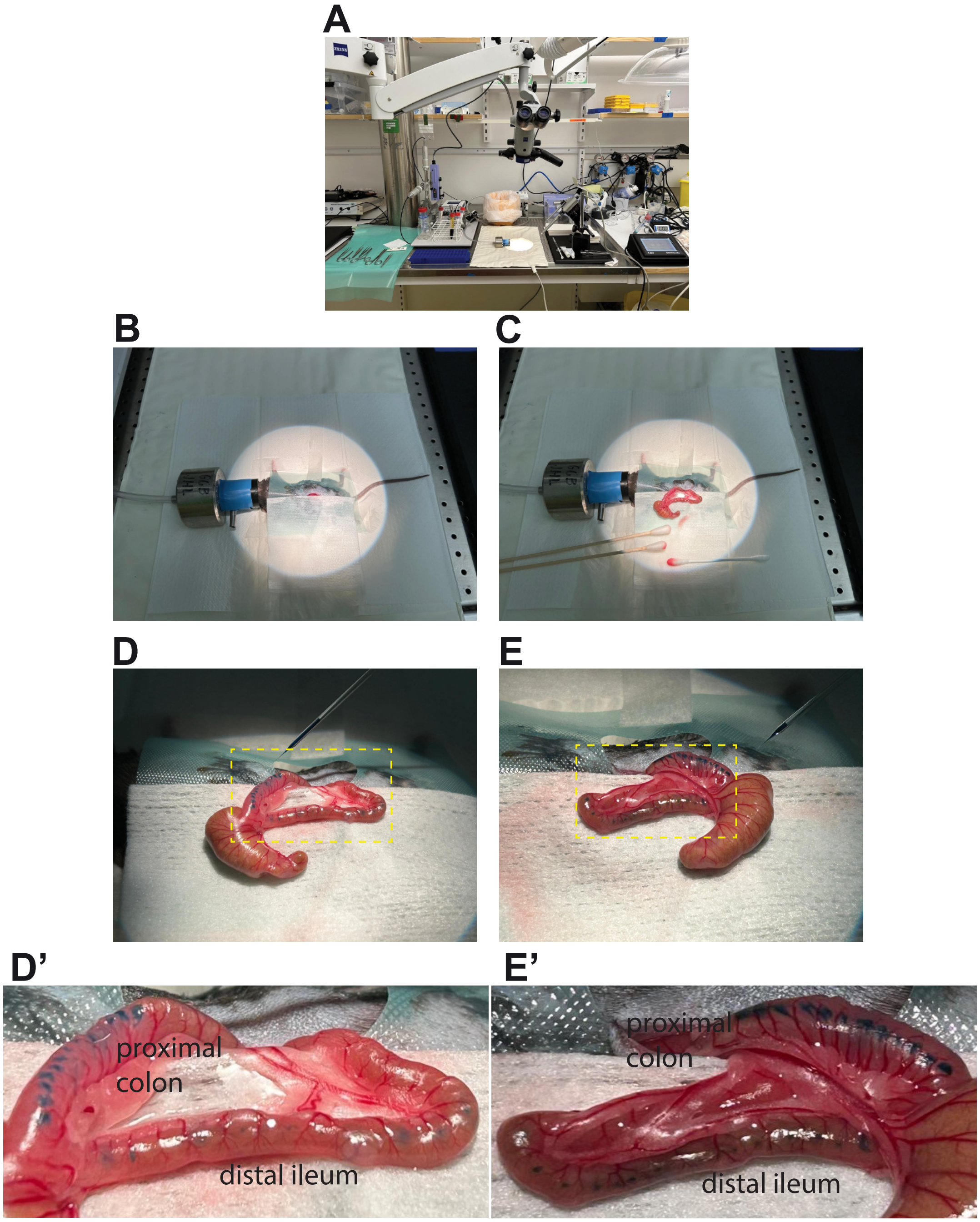

**Supplementary Figure S1. Experimental procedure for local injection into the mouse intestine. (A)** Overview of the surgery setup, showing heated pad, surgical tools and the nanoinjector**. (B)** An incision is made in the center of the abdomen. **(C)** The distal ileum and proximal colon are located, taken out, and placed on a gauze soaked with DPBS. **(D, E)** Injections are made using glass pulled pipettes, 10-15 injection on one side of the distal ileum and another 10-15 injections on the other side. In the case of revealing the circuits between enteric neurons and CG-SMG (Figure 1K, L), an additional 10-15 injections are made in the proximal colon on each side. (**D', E'**) Higher magnification pictures of areas in (D, E) indicated by yellow dotted boxes, focusing on the injection sites in the distal ileum and proximal colon.
