## Supplementary Figure 2 for "Mapping Enteric Neural Circuits by Anterograde Transsynaptic Tracing"

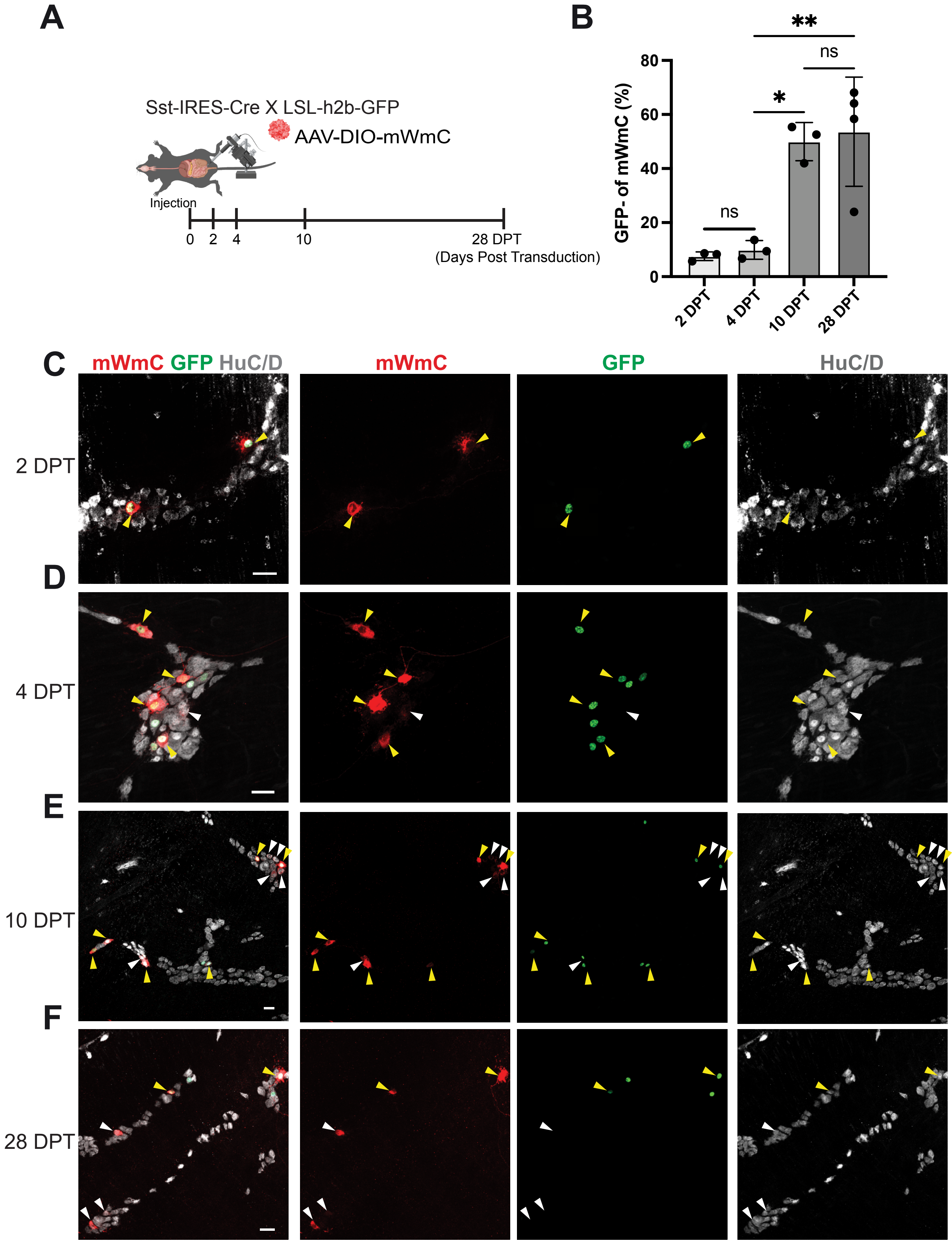

**Supplementary Figure S2. Time course of mWmC transsynaptic tracing in the ENS. (A)** Schematic showing tissue collection days post transduction (DPT) to determine the time-point of mWmC transsynaptic transmission using Sst-IRES-Cre x LSL-h2b-GFP mice. (**B)** Bar graph displaying the number of mWmC+/EGFP- neurons (target neurons) at 2, 4, 10 and 28 DPT. Each dot represents one mouse. N=3-4 mice. Data are presented as mean ± S.D. (**C-F)** Confocal images showing mWmC+ neurons that are either EGFP+ (yellow arrowhead) or EGFP- (white arrowhead). 2 DPT vs 4 DPT, P=0.9952; 4 DPT vs 10 DPT, P=0.0134; 4 DPT vs 28 DPT, P=0.0052; 10 DPT vs 28 DPT, P=0.9782). One-way ANOVA with Tukey's multiple-comparison test were used to determine the statistical significance; *P < 0.05; **P < 0.01; non-significant (ns) P>0.05. Scale bars: 30µm. Part of **(A)** is created with Biorender.com.
